# Targeted degradation of microtubule-associated protein tau using an engineered nanobody-E3 ubiquitin ligase adapter fusion

**DOI:** 10.1101/2022.09.01.506229

**Authors:** Shiyao Wang, Shaowei Jiang, Guoan Zheng, Yong Ku Cho

**Affiliations:** Department of Chemical and Biomolecular Engineering, University of Connecticut, Storrs, CT 06269, United States; Department of Biomedical Engineering, University of Connecticut, Storrs, CT 06269, United States; Institute for Systems Genomics, CT Institute for the Brain and Cognitive Sciences, University of Connecticut, Storrs, CT 06269, United States

**Keywords:** nanobody, protein degradation, microtubule-associated protein tau, SPOP, neuron

## Abstract

Reducing the level of microtubule-associated protein tau has recently emerged as a promising therapeutic approach for a range of neurodegenerative diseases. Among the various approaches, targeted protein degradation provides a reversible means to rapidly reduce and specifically target disease-relevant forms of tau. However, in aging cells, the protein turnover activity is generally weakened, reducing the efficacy of protein degradation. A potential solution to this is to harness the nuclear proteasomal activity. The nucleus has a high proteasomal content and the degradation activity remains relatively unaffected even in aged cells. Here we show that an E3 ligase F-box domain from the nuclear protein human speckle type BTB/POZ protein (SPOP) is effective in degrading the microtubule-associated protein tau in primary mouse hippocampal neurons. Using EGFP-tagged tau and a GFP-binding nanobody fused to SPOP, we found that the native nuclear localization signal in SPOP causes nuclear sequestration of the target protein. However, degradation of the sequestered target proteins is incomplete, resulting in nuclear accumulation. Replacing the native SPOP nuclear localization signal (NLS) with variants having altered nuclear localization efficiency dramatically affects in the degree of nuclear accumulation of the target protein. Interestingly, nanobody-SPOP with no NLS was more efficient than that with a NLS in reducing overall tau level, causing an approximately 50% reduction in ectopically expressed human tau in mouse neurons. These results show the potential for harnessing the nuclear proteasomal activity for targeted tau degradation in cells and demonstrate a new modality of regulating intracellular protein degradation.

## Introduction

Growing evidence shows striking correlations between molecular signatures within microtubule-associated protein tau and neurodegenerative diseases. In Alzheimer’s disease (AD), tau positron emission tomography intensity correlates with atrophy (1) and discriminates patients with high specificity (2). An increase in the plasma tau concentration is associated with AD dementia risk (3, 4), and plasma levels of tau phosphorylated at specific residues emerged as a highly accurate biomarker of AD (5–9). Although these observations show a clear association with tau in neurodegeneration, whether tau causes neurotoxicity and the mechanism of action remain as active areas of investigation (10, 11).

Knocking out or lowering tau expression allows testing of the causality of tau and may lead to novel therapeutic modalities. Although tau stabilizes microtubules and supports cargo transport, neither reduction nor complete knockout of tau alters axonal transport in cultured neurons (12, 13). In rodents, tau knockout is tolerated without apparent histological abnormality in the nervous system (14), particularly during early development (15–17). However, in AD mouse models, tau reduction prevents behavioral deficits (18) and rescues neuronal damage near amyloid deposits (19). In mouse models, tau knockout and reduction also led to discovery of many surprising beneficial effects, including preventing autism (16) and stroke (20) and increased resistance to epilepsy (21, 22).

Although no apparent brain dysfunction is observed in tau knockout mice, a large body of evidence exists on the subtle effects of complete tau knockout on diverse signaling pathways (10, 11). Tau knockout mice show age-dependent obesity potentially due to an impaired hippocampal response to insulin (23, 24), abnormal sleep-wake cycle (25), and impaired adult neurogenesis (26). Several studies reported deficits in motor function (27) and learning and memory (28) in tau knockout mice, although these effects were not consistently observed (24, 29). Due to these potential concerns, strategies for the acute reduction of tau levels are gaining attention. Targeted degradation strategies enable the rapid reduction of cellular proteins and can be implemented with reversible control mechanisms (30–33).

A major concern regarding protein degradation strategies is the impairment of protein turnover in aging cells. Protein degradation by the lysosome and proteasome are both compromised in aged cells (34, 35). This is closely linked to the accumulation of oxidized proteins, unfolded proteins, aggregates, and abnormal post-translational modifications (36, 37). A potential solution to this is to harness the nuclear proteasomal activity. The nucleus has high proteasomal content (38, 39) and can degrade damaged proteins during oxidative stress (40, 41), and the degradation activity remains relatively unaffected in aged cells (42).

To this end, we sought to leverage an E3 ligase F-box domain from a nuclear protein, the human speckle type BTB/POZ protein (SPOP), for targeted tau degradation. SPOP is localized in the nucleus under normal conditions (43) and ubiquitinates proteins in the nucleoplasm (44). When the substrate binding domain of SPOP was replaced with nanobodies, the targeted degradation activity of SPOP was primarily restricted to the nucleus, resulting in low degradation efficiency of cytoplasmic proteins (45, 46). We noted that the endogenous SPOP domain contains a C-terminal nuclear localization signal (NLS) sequence (47); however, the NLS was deleted in previous nanobody-SPOP constructs (45, 46). Since the NLS facilitates the transport of proteins to the nucleus (48), we postulated that the targeted degradation efficiency of cytoplasmic proteins mediated by nanobody-SPOP fusions would be enhanced by restoring the NLS to the SPOP.

We discovered that nanobody-SPOP fusions, including the native C-terminal NLS of SPOP, cause nuclear sequestration of the target protein. However, degradation of the sequestered target proteins is incomplete, resulting in nuclear accumulation. Replacing the native SPOP NLS with variants having altered nuclear localization efficiency dramatically affects in the degree of nuclear accumulation of the target protein. We found that nanobody-SPOP with a NLS derived from the *E. coli* replication fork arresting protein Tus eliminates the nuclear accumulation of EGFP-fused tau. Moreover, interestingly the modified nanobody-SPOP construct selectively reduces tau dissociated from the microtubule. The selective degradation pattern mediated by nanobody-SPOP constructs was consistently observed in primary mouse hippocampal neurons. Unexpectedly, nanobody-SPOP with no NLS was more efficient than that with a NLS in reducing the overall level of tau, decreasing ectopically expressed human tau in mouse neurons by approximately 50%. These results show the potential for harnessing the nuclear proteasomal activity for targeted tau degradation in cells and demonstrate a new modality of regulating intracellular protein degradation.

## Methods

**Table 1.**
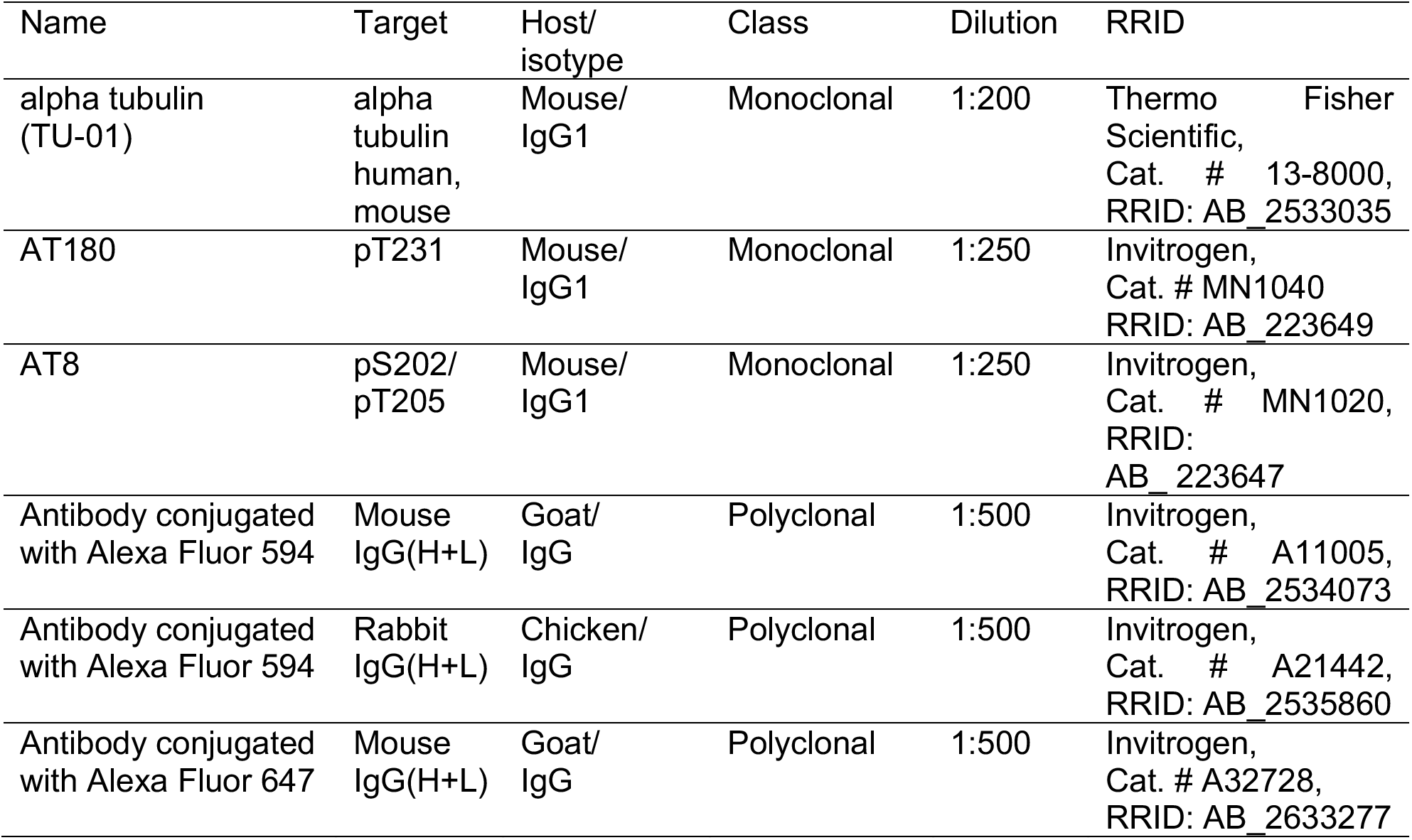
Antibodies

### Cell culture and transfection

Human embryonic kidney (HEK) 293FT cells were purchased from ThermoFisher (# R70007, Research Resource Identifier, RRID: CVCL_ 6911) and grown in Dulbecco’s modified Eagle’s medium (ThermoFisher, # 12320-232, low glucose) supplemented with 10% fetal bovine serum (FBS). To allow confocal microscopy, HEK293FT cells were cultured on Matrigel (Corning Life Sciences, #354234, Tewksbury, MA, USA) coated glass coverslips (Neuvitro, #1.5, acid treated, 12 mm). The glass coverslips were placed in the wells of 24-well plates to aid Matrigel coating and subsequent procedures. The same number of HEK293FT cells were seeded on 2 days before plasmid transfection to reach 70% confluency. The cells were transfected using the Lipofectamine 3000 transfection reagent (Invitrogen, Carlsbad, CA, USA, # L3000001) according to the manufacturer’s instructions.

### Plasmid construction

The nanobody-SPOP constructs were cloned into the pEGFP-N3 vector (Clontech) using restriction enzymes BamHI, SpeI, or BsrGI (New England Biolabs). Sequences of cAbGFP4, LaG2, LaM4, and wild-type SPOP were synthesized by Integrated DNA Technologies (Coralville, IA, USA). SPOP NLS variants were cloned by using flanking primers listed in **Supp. Table 1**. Mammalian expression plasmids pRK5-EGFP-tau and pcDNA3-GSK3b were obtained from Addgene (Watertown, MA, USA). The pRK5-EGFP-tau (# 46904, RRID: Addgene_46904) encodes EGFP fused to human tau isoform 0N4R, which contains four C-terminal repeat regions and lacks the N-terminal insertion sequences. The pcDNA3-GSK3b (# 14754, RRID: Addgene_14754) encodes a hemagglutinin tagged constitutively active mutant (S9A) of glycogen synthase kinase-3β (GSK-3β).

The neuronal expression plasmid Fck-ChR2-GFP (49) (a gift from Ed Boyden) contains a calcium/calmodulin-dependent protein kinase II (CaMKII) alpha promoter that enables heterologous gene expression in hippocampal neurons (50). Full length human tau, which contains four C-terminal repeat regions and two N-terminal sequences (2N4R) was synthesized by Integrated DNA Technologies. The EGFP-tau gene was cloned using overlap extension PCR, and then cloned into the Fck vector using restriction enzymes BamHI and EcoRI (New England Biolabs). Fck-cAbGFP4-SPOP and SPOP NLS variants were cloned by replacing EGFP-tau using restriction sites BamHI and NheI.

### Dissociated mouse hippocampal neuron culture and transfection

The procedures for mouse hippocampal dissection were conducted according to the US National Institutes of Health Guide for the Care and Use of Laboratory Animals and approved by the University of Connecticut Institutional Animal Care and Use Committee (approved protocol A18-004). Hippocampal dissection of postnatal day 0 or day 1 Swiss Webster mice (Taconic) and cell dissociation were performed according to the methods previously described (51, 52). Dissociated cells were seeded on Matrigel coated glass coverslips in 24-well plates and cultured for 2 days in minimum essential medium (Gibco, without phenol red) with 5 g/L glucose, 0.1 g/L bovine transferrin (Sigma), 2.38 g/L HEPES, 2 mM L-glutamine, 0.025 g/L insulin, 10% (v/v) FBS, and B27 supplement (Gibco, diluted to 1x), pH 7.3. Cells were then transfected using a previously described calcium-phosphate transfection method (53). In each transfection, the total amount of plasmid DNA added per well was fixed at 1.25 μg. In wells transfected with tdTomato, EGFP-tau, and cAbGFP4-SPOP NLS variants, 0.42 μg of each plasmid DNA was added. In wells transfected with tdTomato and EGFP-tau, 0.42 μg of a non-expressing control plasmid (pUC18) was added to compensate for the absence of the plasmid containing nanobody-SPOP fusions.

### Immunocytochemistry

HEK293FT cells were seeded on Matrigel coated glass coverslips in 24-well plates, then co-transfected with plasmids pRK5-EGFP-tau and an empty vector pUC-18 (ThermoFisher) to equalize the amount of plasmid DNA, or co-transfected with pRK5-EGFP-tau and pcDNA3-GSK-3β S9A. After 16-20 hours of expression, cells were washed with ice-cold PBSCM (37 mM NaCl, 2.7 mM KCl, 10 mM Na_2_HPO_4_, 1.8 mM KH_2_PO_4_, 1 mM CaCl_2_, and 0.5 mM MgCl_2_, pH 7.4), then fixed in 4% paraformaldehyde (dissolved in PBSCM, pH 7.4) for 15 min at 25°C, and then washed three times with ice-cold PBSCM. The fixed cells were permeabilized using PBSCM with 0.1% Triton X-100 for 15 min at 25°C, then washed three times with ice-cold PBSCM. Samples requiring dephosphorylation were treated with 2 U/μL λPP (New England Biolabs, # P0753) for 3 h at 30°C and then washed with PBSCM. After the permeabilization and dephosphorylation treatment, cells were blocked in PBSG (PBSCM with 40% goat serum; Gibco, # 16210-064) for 30 min at 25°C and then incubated with primary antibodies (dilution listed in **Table 1**) diluted in PBSG overnight at 4°C in the dark. The next day, cells were washed three times with ice-cold PBSCM (5 min incubation on ice in between) and then incubated for 2 hours at 4°C in the dark with Alexa Fluor 647 conjugated secondary anti-mouse antibody, Alexa Fluor 594 conjugated secondary anti-mouse antibody, or Alexa Fluor 594 conjugated secondary anti-rabbit antibody (dilutions listed in **Table 1**) diluted in PBSG. Cells were then washed three times with ice-cold PBSCM (5 min incubation on ice in between) before the sample preparation for confocal imaging.

### Sample preparation and confocal imaging

After plasmid expression, cells were washed with ice-cold PBSCM, fixed in 4% paraformaldehyde (dissolved in PBSCM, pH 7.4) for 15 min at 25°C, and washed three times with ice-cold PBSCM. Immunocytochemistry was conducted according to the previous section for cells requiring antibody staining. The glass coverslips on which cells grown on were picked up using tweezers and mounted on glass slides with ProLong™ Diamond antifade mountant (Thermofisher, # P36965), which contains 4’,6-diamidino-2-phenylindole (DAPI) that stains the nuclei. After 24 hours of curing at 25°C in the dark, cells were imaged using a Nikon A1R confocal microscope (UConn Center for Open Research Resources and Equipment).

### Colocalization and protein distribution analysis in neurons

Quantitative analysis of colocalization and protein distribution was performed using MATLAB (MathWorks). The first step of the analysis was to define the region of interest (ROI) to cut out the region containing EGFP-tau-expressing cells. For this, the tdTomato channel was binarized by setting a threshold value based on the image intensity histogram. Considering the co-transfection of EGFP-tau and tdTomato, the ROI mask was generated based on the tdTomato-expressing cell region. The ROI can be extracted from these images by multiplying the ROI mask with the tdTomato channel, EGFP-tau channel, and microtubule channel. Only the ROI of each channel was used for all the following colocalization analyses and fluorescence quantifications.

The ratio of the total amount of EGFP-tau to tdTomato was quantified using the following equation:

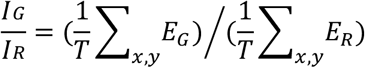

where 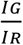 is the ratio, *E_G_* and *E_R_* represent the pixel intensity in the ROIs of the EGFP-tau and tdTomato channels, respectively, and *T* is the total number of pixels in the ROI.

Segmentation of the nuclear and cytosolic regions was conducted using the following three steps to quantify the intracellular distribution of EGFP-tau. First, an intensity map was generated by dividing the tdTomato channel by the microtubule channel to enhance the contrast:

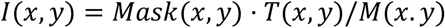

where *I*(*x, y*) is the intensity map after processing, *Mask*(*x, y*) is the ROI mask, “·” stands for point-wise multiplication, and *T*(*x, y*) and *M*(*x.y*) are the raw images of the tdTomato channel and microtubule channel, respectively. Second, *I*(*x, y*) was binarized to extract the nuclear region using Otsu’s method (54) by minimizing the intraclass variance of the thresholded pixels. Third, segmentation was performed using morphological opening and closing functions in MATLAB and manual refining to remove other small regions. The cell nucleus and cytosol of the EGFP-tau channel were then separated based on the segmentation.

The intracellular distribution of EGFP-tau was quantified using the following equation.

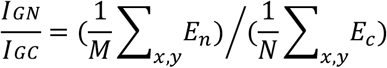

Here, 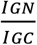 is the distribution coefficient, *E_n_* and *E_c_* represent the pixel intensity in the nuclear and cytosolic regions of the EGFP-tau channel, and *M* and *N* are the numbers of pixels of *E_n_* and *E_c_* in the segmentation, respectively.

The colocalization of EGFP-tau and microtubule was analyzed by plotting the pixel intensity in the image which reveals the intensity correlation between the two proteins (with respect to the protein distribution in the nuclear and cytosolic regions using red and blue colors, respectively).

## Results

### Nanobody-SPOP fusion causes nuclear accumulation of target proteins due to the native SPOP nuclear localization signal

SPOP contains a substrate-binding MATH (meprin and traf homology) domain; a BTB (bric à brac, tramtrack, broad complex) dimerization domain; another dimerization domain, BACK (BTB and C-terminal Kelch), which contributes to tetramer formation; and a C-terminal NLS sequence (47, 55, 56). In previous studies, the NLS of the SPOP domain (**Fig. 1a**) was deleted in nanobody-SPOP fusion constructs. We hypothesized that a nanobody-SPOP construct with its native SPOP NLS would lead to nuclear sequestration of the target protein, resulting in the degradation of cytoplasmic proteins. To test this, we first conducted targeted degradation of the enhanced green fluorescent protein (EGFP), which localizes throughout the cell when expressed in the human embryonic kidney 293FT (HEK293FT) cell (**Fig. 1b**). Consistent with previous reports (45, 46), cAbGFP4 fused to SPOP without the NLS (cAbGFP4-SPOPΔNLS) resulted in efficient degradation of nuclear EGFP but not cytoplasmic EGFP (**Fig. 1b**). When we co-expressed the EGFP and a nanobody binding to EGFP (cAbGFP4) (57) fused with SPOP, including its native NLS, we observed nuclear sequestration of EGFP (**Fig. 1b**). When we replaced cAbGFP4 with another EGFP nanobody (LaG2) targeting a different epitope (58, 59), EGFP was again localized in the nucleus, although the pattern of nuclear accumulation differed (**Fig. 1b**). To assess whether the degradation pattern was specific to EGFP, we conducted analogous experiments using a mCherry targeting nanobody (LaM4) (58). As with EGFP, while LaM4-SPOPΔNLS caused degradation of nuclear mCherry, LaM4-SPOP resulted in nuclear accumulation of mCherry (**Fig. 1c**). These results indicate that the SPOP NLS causes nuclear sequestration of target proteins of nanobody-SPOP fusions but results in incomplete degradation.

**Figure 1.**
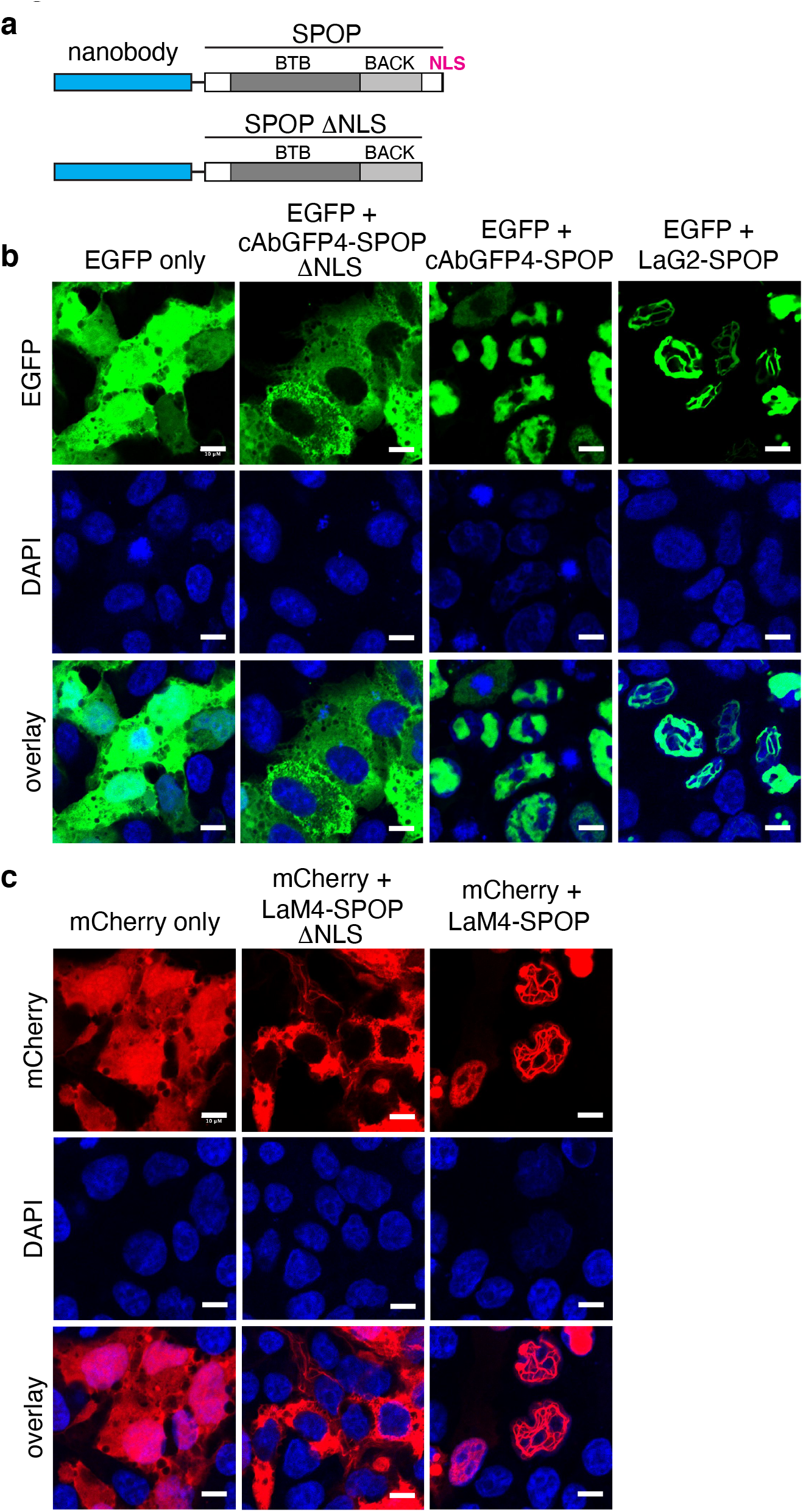
A native SPOP NLS causes nuclear accumulation of proteins targeted by nanobody-SPOP fusions. **(a)** A schematic summary of nanobody-SPOP fusion constructs used. Nanobodies are genetically fused to the full-length SPOP or SPOP without the NLS (SPOPΔNLS). **(b, c)** Confocal microscopy images of HEK293FT cells transfected with fluorescent proteins EGFP (b) or mCherry (c) with or without the indicated nanobody-SPOP constructs. Nanobodies cAbGFP4 and LaG2 bind to GFP, while LaM4 binds to mCherry. Scale bars indicate 10 μm.

### Altering the NLS sequence fused to nanobody-SPOP prevents nuclear accumulation and enables visualization of microtubule-bound tau

Depending on its sequence, the NLS causes varying degrees of enrichment of fused proteins in the nucleus (60). We speculated that the accumulation of EGFP in the nucleus is caused by the NLS of SPOP (**Fig. 2**). We hypothesized that altering the NLS of SPOP may enable the degradation of cytoplasmic proteins without causing nuclear accumulation. Although the microtubule-associated protein tau primarily localizes in the cytoplasm (**Fig. 3a**), we anticipated that targeting it with a nanobody-SPOP fusion would result in nuclear accumulation, as observed when targeting EGFP. Indeed, when we co-expressed EGFP-fused human 0N4R tau (EGFP-tau, EGFP fused to the N-terminus of tau) and cAbGFP4-SPOP in HEK293FT cells, EGFP-tau showed apparent nuclear localization (**Fig. 3a**). We substituted the native NLS of SPOP with four distinct NLS sequences, derived from the simian virus 40 large T antigen (SV40) (61), c-Myc (62), nucleoplasmin (NLP) (63), or *E. coli* replication fork arresting protein Tus (Tus) (64) (**Fig. 3b**). The cAbGFP4-SPOP with an altered NLS resulted in dramatic changes in the localization pattern of EGFP-tau (**Fig. 3c**). Confocal microscopy confirmed the nuclear accumulation of EGFP-tau when targeted with cAbGFP4-SPOP, but not with LaM4-SPOP (**Fig. 3d**). Notably, nuclear accumulation of EGFP-tau was absent in cells co-transfected with cAbGFP4-SPOP fused with the NLS from Tus (cAbGFP4-SPOP-Tus) (**Fig. 3d**). In these cells, the localization pattern of EGFP-tau was reminiscent of microtubules, and indeed co-localized with the microtubule component a-tubulin (**Fig. 3d**). EGFP-tau also showed similar co-localization when co-expressed with cAbGFP4-SPOPΔNLS (**Fig. 3d**), although the localization pattern was less clear than that observed with cAbGFP4-SPOP-Tus. These results suggest that modifying the NLS prevented nuclear accumulation of EGFP-tau, probably by reducing the nuclear import rate; however, a significant fraction of EGFP-tau bound to microtubules remained. When we targeted EGFP using cAbGFP4-SPOP-Tus, punctate nuclear EGFP was observed, indicating that cAbGFP4-SPOP-Tus causes nuclear import (**Fig. 3d**). These results indicate that targeting EGFP-tau with cAbGFP4-SPOP-Tus preferentially reduces unbound EGFP-tau but does not significantly reduce the overall level of EGFP-tau in transiently transfected HEK293FT cells.

**Figure 2.**
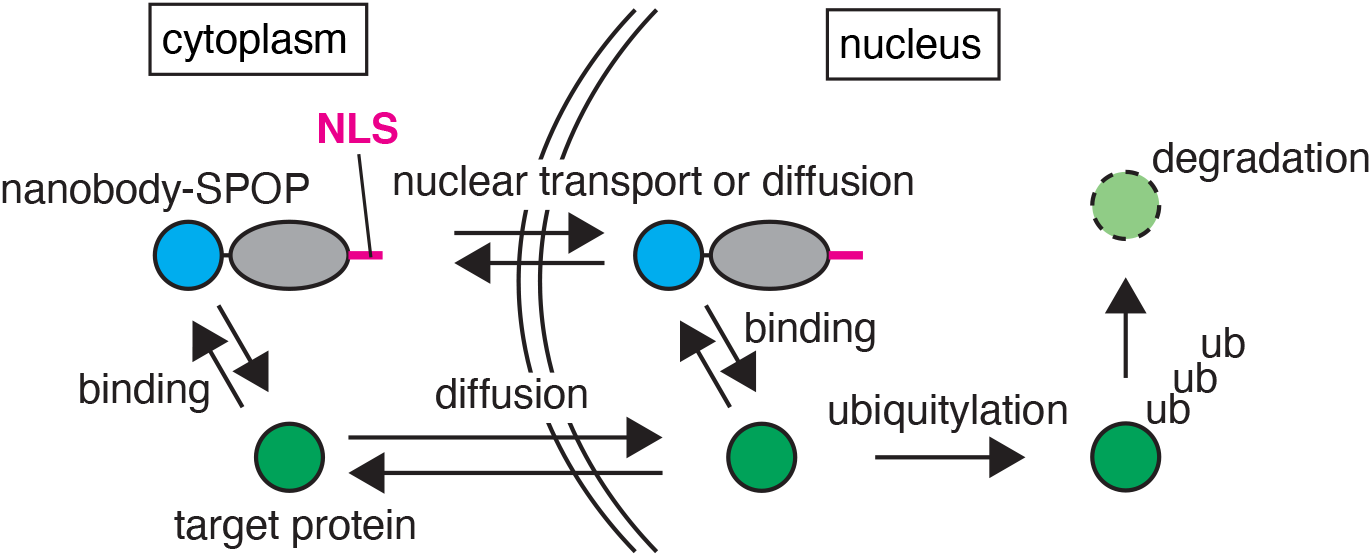
A summary of processes in nuclear degradation mediated by nanobody-SPOP. Blue indicates nanobody, grey shade indicates SPOP, and green indicates the target protein. The NLS is indicated by magenta.

**Figure 3.**
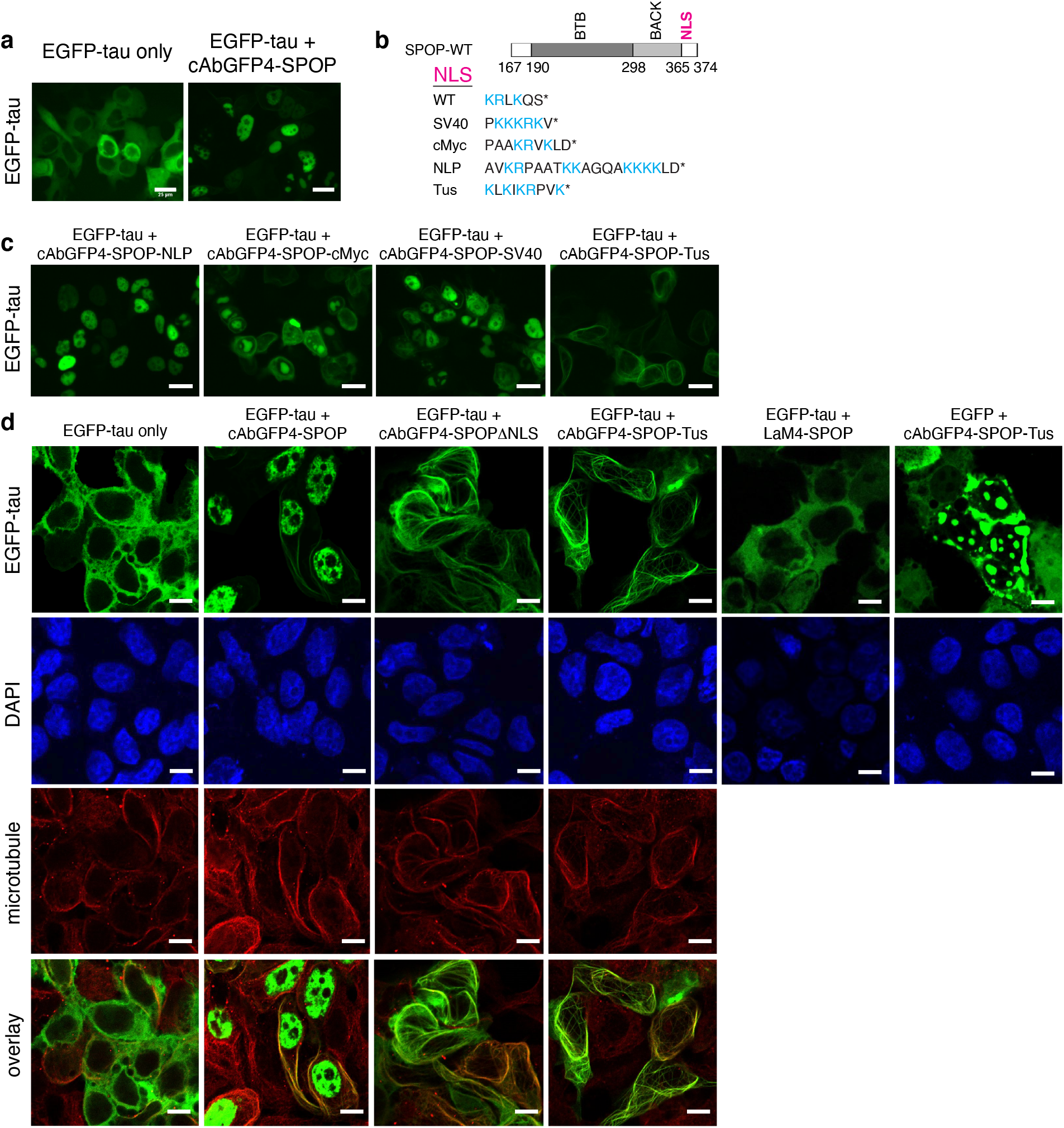
Targeting EGFP-tau using nanobody-SPOP with varying NLSs. **(a)** Fluorescence microscopy images of HEK293FT cells transfected with EGFP-tau with or without cAbGFP4-SPOP. **(b)** A summary of NLS variants used. **(c)** Fluorescence microscopy images of HEK293FT cells transfected with EGFP-tau with cAbGFP4-SPOP NLS variants. Scale bars in panels (a) and (c) indicate 25 μm. **(d)** Confocal microscopy images of HEK293FT cells transfected with the indicated constructs labeled with an anti-a-tubulin antibody. Scale bars indicate 10 μm.

### Imaging distinctive phosphorylation patterns in microtubule-bound fraction of tau

Targeting EGFP-tau with cAbGFP4-SPOP-Tus opens new possibilities for assessing microtubule-bound tau in cells. Based on previous *in vitro* studies suggesting that tau phosphorylation impacts its microtubule binding, we hypothesized that the microtubule-bound fraction of tau revealed after targeting with cAbGFP4-SPOP-Tus will have distinct phosphorylation patterns. After co-expressing EGFP-tau and cAbGFP4-SPOP-Tus, we fixed the cells and stained them with previously validated high specificity phosphorylated tau (p-tau) antibodies (52). HEK293FT cells transfected with EGFP-tau showed clear staining with AT180 (pT231 tau) and AT8 (pS202/pT205 tau) (**Fig. 4**). We observed a clear enhancement of AT8 staining when an activated form of glycogen synthase kinase 3β (GSK-3β) was co-expressed, as we reported previously (52). In both cases, treatment of cells using lambda phosphatase eliminated antibody staining (**Fig. 4**). In cells co-expressing EGFP-tau and cAbGFP4-SPOP-Tus, AT180 staining was detected, but not for AT8 even under overexpression of GSK-3β (**Fig. 4**). This result shows that the microtubule-bound fraction of tau is phosphorylated at T231 but not at S202 and T205, while the total tau is phosphorylated at all these residues.

**Figure 4.**
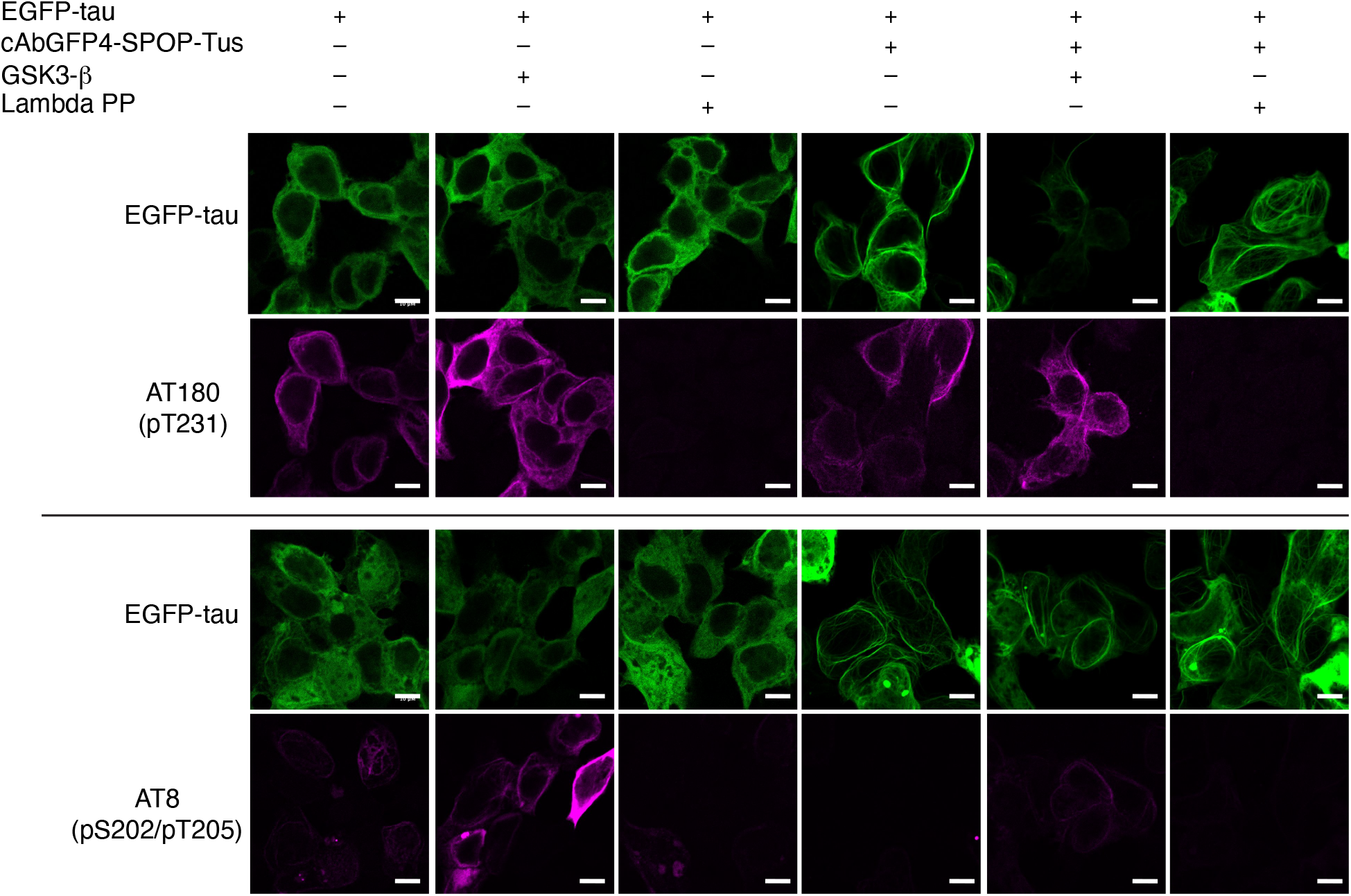
Distinct phosphorylation patterns of microtubule-bound tau in HEK293FT cells were revealed using targeted degradation. Cells were transfected with EGFP-tau with or without cAbGFP4-SPOP-Tus. In some samples, GSK-3β was also co-transfected to enhance tau phosphorylation. Some samples were treated with lambda protein phosphatase to dephosphorylate tau. The cells were labeled with monoclonal p-tau antibodies AT180 (specific to tau phosphorylated at threonine 231) or AT8 (specific to tau phosphorylated at serine 202 and threonine 205). Scale bars indicate 10 μm.

### Engineered nanobody-SPOP significantly reduces EGFP-tau mouse primary neurons without causing nuclear accumulation

Since the microtubule-associated protein tau is primarily expressed in neurons, we tested targeted degradation using the engineered nanobody-SPOP constructs in primary neuron cultures. We expressed human 2N4R tau as an EGFP fusion (EGFP-tau) in mouse primary hippocampal neurons (**Fig. 5a**) using a CaMKII alpha promoter. This promoter is highly active in hippocampal neurons (50) and has been used to express P301L tau in a mouse model of human tauopathy (rTg4510) (65). As an internal transfection control, a red fluorescent protein (tdTomato) was co-expressed under the same promoter (**Fig. 5a**). Strong EGFP-tau fluorescence was detected in the soma and dendrites, as in previous studies observing ectopically expressed tau (66, 67), and also in the nucleus (**Fig. 5a**). When cAbGFP4-SPOP or cAbGFP4-SPOPΔNLS was co-expressed with EGFP-tau, we observed a significant reduction in EGFP-tau levels normalized to that of tdTomato (**Fig. 5a, b**, 23% and 51% reduction, respectively), but not for cAbGFP4-SPOP-Tus (**Fig. 5a, b**). The cAbGFP4-SPOP co-expression with EGFP-tau resulted in a 3.3-fold increase in nuclear EGFP-tau fluorescence normalized to cytoplasmic EGFP-tau fluorescence (**Fig. 5c, d**), resembling the results from HEK293FT cells (**Fig. 3**). Although cAbGFP4-SPOP-Tus did not significantly lower the level of tau, it reduced the nuclear tau normalized to cytoplasmic tau by 47% (**Fig. 5c**) and resulted in a clear positive correlation between EGFP-tau fluorescence and microtubule immunostaining (**Fig. 5e**). Taken together, these results indicate that in primary neurons, cAbGFP4-SPOPΔNLS significantly reduces tau dissociated from microtubules without causing nuclear accumulation (**Fig. 6**).

**Figure 5.**
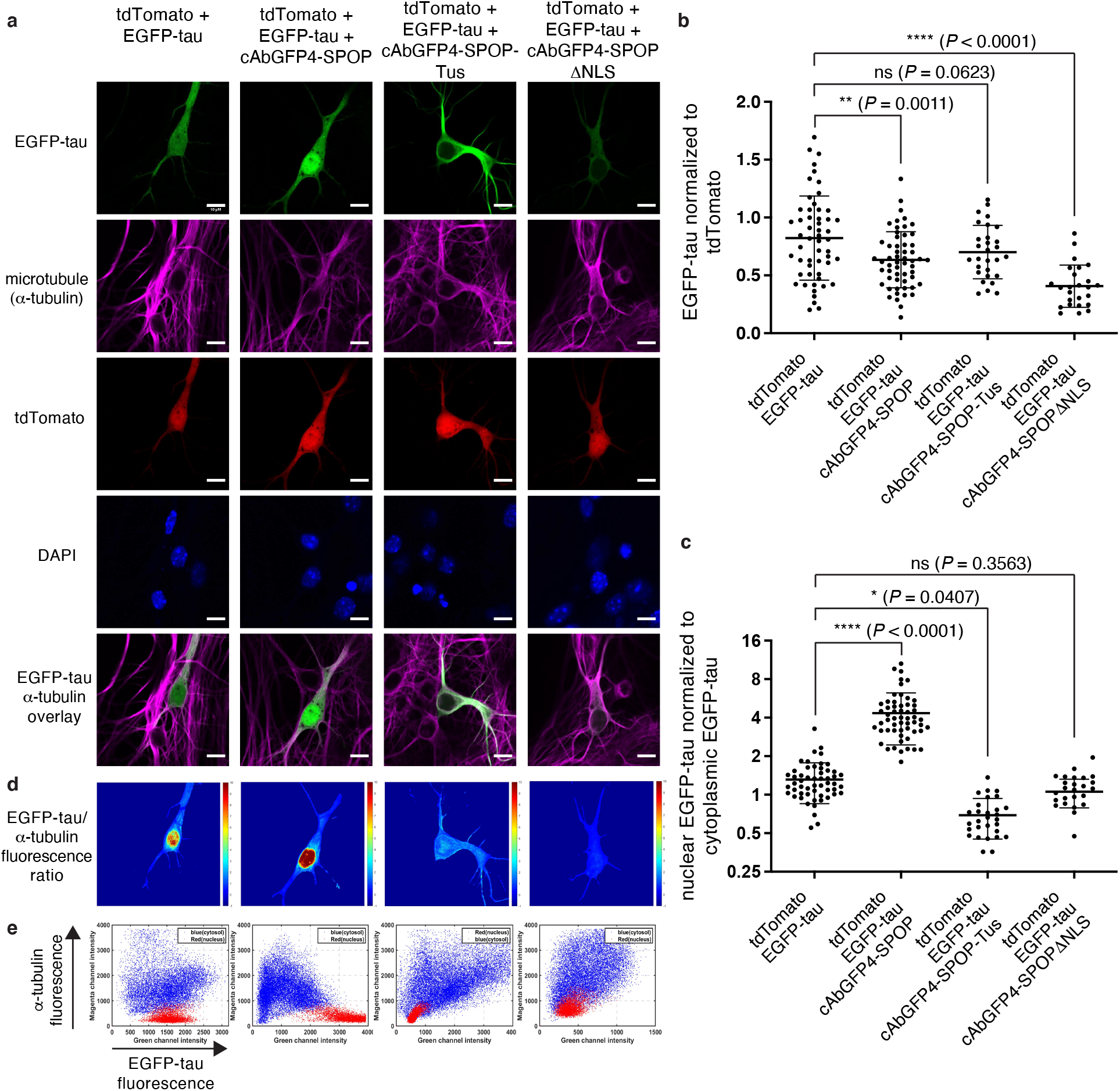
Targeted degradation of EGFP-tau in primary mouse hippocampal neurons. **(a)** Confocal microscopy images of cultured neurons were transfected with tdTomato (transfection control), EGFP-tau, with or without the nanobody-SPOP variants indicated. The cells were also labeled with an anti-a-tubulin antibody. Scale bars indicate 10 μm. **(b)** Comparison of EGFP-tau levels normalized to the tdTomato level. **(c)** Comparison of nuclear EGFP-tau levels normalized to the cytoplasmic EGFP-tau level. **(d)** Heat map images of EGFP-tau levels normalized to the microtubule level. **(e)** Correlation of EGFP-tau fluorescence and a-tubulin fluorescence. In panel (e), red indicates pixels inside the nucleus while blue indicates pixels outside the nucleus. For panels (b) and (c), each datapoint plotted was calculated from a neuron and 25-54 neurons were imaged for each condition across multiple experiments. The P-values were calculated using the Holm-Šídák multiple-comparisons test; *, *P* ≤ 0.05; **, *P* ≤ 0.01; ****, *P* ≤ 0.0001; ns, *P* > 0.05.

**Figure 6.**
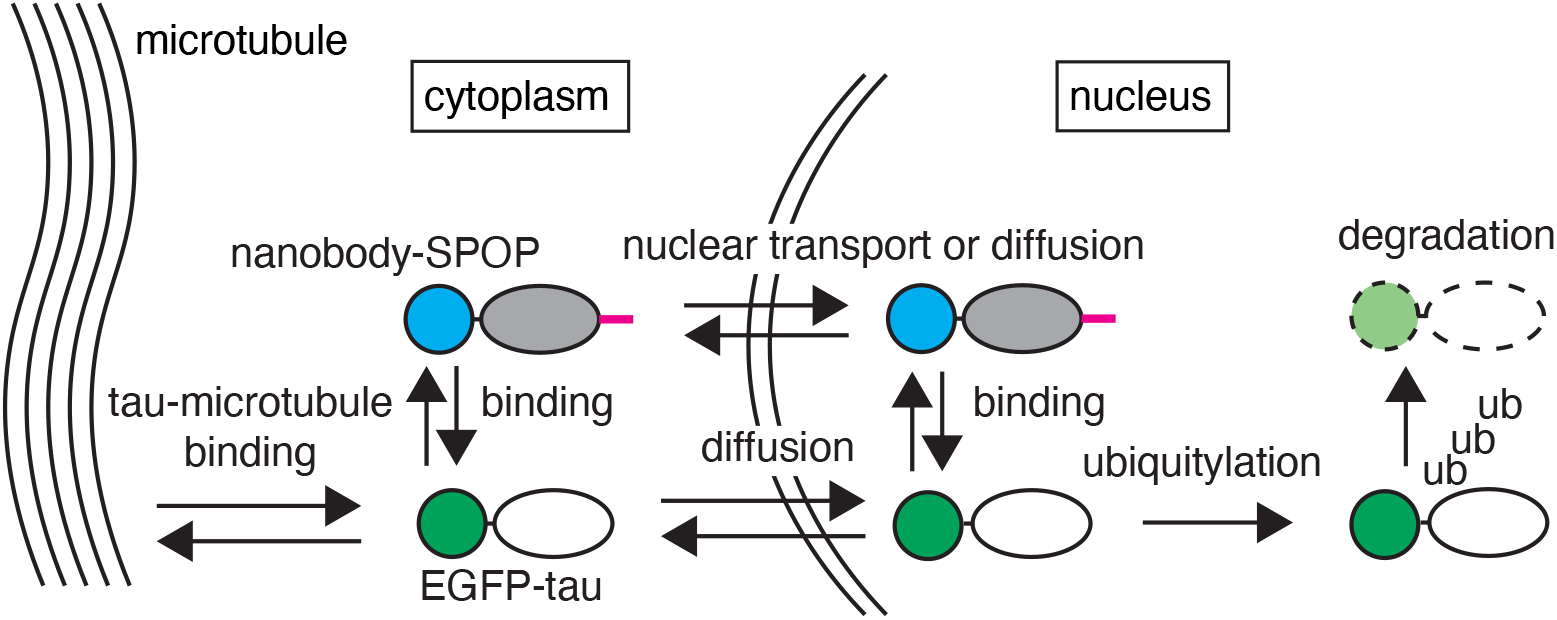
A summary of factors affecting EGFP-tau localization upon targeted degradation using nanobody-SPOP constructs. Blue indicates nanobody, grey shade indicates SPOP, green indicates EGFP, and unfilled oval indicates tau. The NLS is indicated by magenta.

## Discussion

In this study, we demonstrated the use of the nuclear E3 ligase adapter SPOP for degrading the human tau protein. Several strategies have shown promise for the acute reduction of tau levels. Antisense oligonucleotides (ASOs) or small interfering RNA (siRNA) targeting tau have been developed to degrade tau mRNA (68–70). ASOs enabled up to 70% tau reduction in adult mice, lowering tau levels in the brain, interstitial fluid, and the cerebrospinal fluid (68). Tau ASOs reversed preexisting tau aggregates, prevented hippocampal atrophy and extended survival in a mouse model of AD, and reduced tau levels in non-human primates (69). Although ASOs are generally effective in reducing tau levels, since protein post-translational modifications and conformational change are strongly associated with tau pathology, protein targeting may enable selective degradation of toxic tau species. For degrading tau proteins, proteolysis-targeting chimeras (PROTACs) using a tau positron emission tomography probe as the targeting moiety reduced the total tau in neurons by approximately 60% (71). Tau-binding peptides fused to E3 ligase-binding peptides have been reported (72, 73); however, their efficiency in neurons has not been determined. Rather than using small molecules for targeting, a single-chain antibody fragment (scFv) targeting tau was also applied in targeting intracellular tau for degradation (74). In this approach, the scFv was directly fused to ubiquitin, which can be polyubiquitylated and recruit the bound tau for proteasomal degradation, resulting in an approximately 30% reduction of tau in neurons (74). In our study, we observed approximately 50% reduction of tau using the nanobody-SPOPΔNLS in primary mouse neurons. Although the degradation efficiency did not differ significantly from previous results, a unique feature of our approach is using the nuclear proteasome to degrade the tau protein. The potential for harnessing the nuclear proteasomal degradation activity in aging cells remains to be tested.

Interestingly, we found that the cAbGFP4-SPOP-Tus mediates the selective reduction of a microtubule-dissociated fraction of tau. This allowed us to preferentially detect the tau fraction bound to the microtubule and probe its phosphorylation status. Our data show selective reduction of tau phosphorylated at S202/T205, compared to tau phosphorylated at T231. Although it is generally accepted that hyperphosphorylation of tau reduces its binding to microtubules (75, 76), the enrichment of a certain phospho-epitope in the microtubule-unbound fraction does not necessarily indicate that these phosphorylations affect microtubule binding. This is because intracellular tau is generally phosphorylated at multiple sites due to endogenous kinases, even when active GSK-3β is not overexpressed (52). Previous studies have found enrichment of pS202/pT205 tau as well as pT231 tau in the microtubule-unbound fraction of cells (76–79); however, a fraction of pT231 tau colocalizes to microtubules (80). Our data corroborate these results, and indicate that in HEK293FT cells, pS202/pT205 tau is more enriched in the microtubule-unbound fraction of tau, compared to pT231 tau. It is noteworthy that tau dissociation from microtubules is thought to contribute to increased cytoplasmic concentration and the formation of toxic oligomers (81). Therefore, reducing the microtubule-unbound fraction of tau may be effective in preventing toxic tau oligomer formation. Moreover, the ability to visualize microtubule-bound tau will be useful in observing the dynamics of tau-microtubule interaction in live cells.

Further investigation is required to determine why cAbGFP4-SPOPΔNLS is more efficient than cAbGFP4-SPOP-Tus in lowering EGFP-tau levels. Since the cAbGFP4-SPOPΔNLS protein does not contain a NLS for active nuclear import, it must primarily rely on passive diffusion to reach the nucleus. Therefore, the primary driving force for the overall reduction of tau is continuous diffusion due to the concentration gradient generated by depletion in the nucleus. When EGFP-tau and cAbGFP4-SPOP-Tus are co-expressed, the EGFP-tau concentration in the cytoplasm reaches a steady-state level much higher than that in the nucleus. This suggests that the diffusion of EGFP-tau to the nucleus is reduced in cells expressing cAbGFP4-SPOP-Tus. It is known that tau binding to certain nuclear pore complex components generates a diffusion barrier (82). Interestingly, this trend was not observed with cAbGFP4-SPOP, as indicated by the increased nuclear concentration of EGFP-tau. This suggests that the diffusion barrier formation depends on the NLS and its partnering importin. Systematic comparison of NLS sequences in neurons may elucidate how the diffusion barrier arises and lead to improvements in the nuclear degradation efficiency.

In summary, we demonstrated that cAbGFP4-SPOPΔNLS enables targeted degradation of EGFP-tau in primary mouse hippocampal neurons. Our data also highlight the importance of nuclear transport and sequestration of proteins in harnessing the nuclear proteasomal activity. This new capability may prove more effective than relying on non-selective proteasomal activity in degrading tau in aging cells, providing a promising modality for reducing tau levels in neurodegenerative diseases.

## Acknowledgements

This work was supported by the NIH grant 1R21NS111358, and Alzheimer’s Association grant 2019-AARG-NFT-640971.

## References

1. La Joie, R., A.V. Visani, S.L. Baker, J.A. Brown, V. Bourakova, J. Cha, K. Chaudhary, L. Edwards, L. Iaccarino, M. Janabi, O.H. Lesman-Segev, Z.A. Miller, D.C. Perry, J.P. O’Neil, J. Pham, J.C. Rojas, H.J. Rosen, W.W. Seeley, R.M. Tsai, B.L. Miller, W.J. Jagust, and G.D. Rabinovici, Prospective longitudinal atrophy in Alzheimer’s disease correlates with the intensity and topography of baseline tau-PET. Science Translational Medicine, 2020. 12(524): p. eaau5732.

2. Leuzy, A., R. Smith, R. Ossenkoppele, A. Santillo, E. Borroni, G. Klein, T. Ohlsson, J. Jögi, S. Palmqvist, N. Mattsson-Carlgren, O. Strandberg, E. Stomrud, and O. Hansson, Diagnostic Performance of RO948 F 18 Tau Positron Emission Tomography in the Differentiation of Alzheimer Disease From Other Neurodegenerative Disorders. JAMA Neurology, 2020. 77(8): p. 955–965.

3. Pase, M.P., A.S. Beiser, J.J. Himali, C.L. Satizabal, H.J. Aparicio, C. DeCarli, G. Chêne, C. Dufouil, and S. Seshadri, Assessment of Plasma Total Tau Level as a Predictive Biomarker for Dementia and Related Endophenotypes. JAMA Neurology, 2019. 76(5): p. 598–606.

4. Mattsson, N., H. Zetterberg, S. Janelidze, P.S. Insel, U. Andreasson, E. Stomrud, S. Palmqvist, D. Baker, C.A. Tan Hehir, A. Jeromin, D. Hanlon, L. Song, L.M. Shaw, J.Q. Trojanowski, M.W. Weiner, O. Hansson, and K. Blennow, Plasma tau in Alzheimer disease. Neurology, 2016. 87(17): p. 1827–1835.

5. Janelidze, S., N. Mattsson, S. Palmqvist, R. Smith, T.G. Beach, G.E. Serrano, X. Chai, N.K. Proctor, U. Eichenlaub, H. Zetterberg, K. Blennow, E.M. Reiman, E. Stomrud, J.L. Dage, and O. Hansson, Plasma P-tau181 in Alzheimer’s disease: relationship to other biomarkers, differential diagnosis, neuropathology and longitudinal progression to Alzheimer’s dementia. Nat Med, 2020. 26(3): p. 379–386. PubMed PMID: 32123385.

6. Mattsson-Carlgren, N., S. Janelidze, S. Palmqvist, N. Cullen, A.L. Svenningsson, O. Strandberg, D. Mengel, D.M. Walsh, E. Stomrud, J.L. Dage, and O. Hansson, Longitudinal plasma p-tau217 is increased in early stages of Alzheimer’s disease. Brain, 2020. 143(11): p. 3234–3241. PubMed PMID: 33068398; PMCID: PMC7719022.

7. Ashton, N.J., T.A. Pascoal, T.K. Karikari, A.L. Benedet, J. Lantero-Rodriguez, G. Brinkmalm, A. Snellman, M. Scholl, C. Troakes, A. Hye, S. Gauthier, E. Vanmechelen, H. Zetterberg, P. Rosa-Neto, and K. Blennow, Plasma p-tau231: a new biomarker for incipient Alzheimer’s disease pathology. Acta Neuropathol, 2021. 141(5): p. 709–724. PubMed PMID: 33585983.

8. Karikari, T.K., T.A. Pascoal, N.J. Ashton, S. Janelidze, A.L. Benedet, J.L. Rodriguez, M. Chamoun, M. Savard, M.S. Kang, J. Therriault, M. Schöll, G. Massarweh, J.-P. Soucy, K. Höglund, G. Brinkmalm, N. Mattsson, S. Palmqvist, S. Gauthier, E. Stomrud, H. Zetterberg, O. Hansson, P. Rosa-Neto, and K. Blennow, Blood phosphorylated tau 181 as a biomarker for Alzheimer’s disease: a diagnostic performance and prediction modelling study using data from four prospective cohorts. The Lancet Neurology, 2020. 19(5): p. 422–433.

9. Arbaciauskaite, M., Y. Lei, and Y.K. Cho, High-specificity antibodies and detection methods for quantifying phosphorylated tau from clinical samples. Antibody Therapeutics, 2021. 4(1): p. 34–44.

10. Chang, C.W., E. Shao, and L. Mucke, Tau: Enabler of diverse brain disorders and target of rapidly evolving therapeutic strategies. Science, 2021. 371(6532). PubMed PMID: 33632820; PMCID: PMC8118650.

11. Wang, Y. and E. Mandelkow, Tau in physiology and pathology. Nat Rev Neurosci, 2016. 17(1): p. 22–35. PubMed PMID: 26631930.

12. Vossel, K.A., J.C. Xu, V. Fomenko, T. Miyamoto, E. Suberbielle, J.A. Knox, K. Ho, D.H. Kim, G.-Q. Yu, and L. Mucke, Tau reduction prevents Aβ-induced axonal transport deficits by blocking activation of GSK3β. Journal of Cell Biology, 2015. 209(3): p. 419–433.

13. Yuan, A., A. Kumar, C. Peterhoff, K. Duff, and R.A. Nixon, Axonal Transport Rates *In Vivo* Are Unaffected by Tau Deletion or Overexpression in Mice. The Journal of Neuroscience, 2008. 28(7): p. 1682–1687.

14. Harada, A., K. Oguchi, S. Okabe, J. Kuno, S. Terada, T. Ohshima, R. Sato-Yoshitake, Y. Takei, T. Noda, and N. Hirokawa, Altered microtubule organization in small-calibre axons of mice lacking tau protein. Nature, 1994. 369(6480): p. 488–491.

15. Morris, M., P. Hamto, A. Adame, N. Devidze, E. Masliah, and L. Mucke, Age-appropriate cognition and subtle dopamine-independent motor deficits in aged Tau knockout mice. Neurobiology of Aging, 2013. 34(6): p. 1523–1529.

16. Tai, C., C.-W. Chang, G.-Q. Yu, I. Lopez, X. Yu, X. Wang, W. Guo, and L. Mucke, Tau Reduction Prevents Key Features of Autism in Mouse Models. Neuron, 2020. 106(3): p. 421–437.e11.

17. Ke, Y.D., A.K. Suchowerska, J. van der Hoven, D.M. De Silva, C.W. Wu, J. van Eersel, A. Ittner, and L.M. Ittner, Lessons from Tau-Deficient Mice. International Journal of Alzheimer’s Disease, 2012. 2012: p. 873270.

18. Roberson, E.D., K. Scearce-Levie, J.J. Palop, F. Yan, I.H. Cheng, T. Wu, H. Gerstein, G.-Q. Yu, and L. Mucke, Reducing Endogenous Tau Ameliorates Amyloid β-Induced Deficits in an Alzheimer’s Disease Mouse Model. Science, 2007. 316(5825): p. 750–754.

19. Wegmann, S., S.L. DeVos, B. Zeitler, K. Marlen, R.E. Bennett, M. Perez-Rando, D. MacKenzie, Q. Yu, C. Commins, R.N. Bannon, B.T. Corjuc, A. Chase, L. Diez, H.B. Nguyen, S. Hinkley, L. Zhang, A. Goodwin, A. Ledeboer, S. Lam, I. Ankoudinova, H. Tran, N. Scarlott, R. Amora, R. Surosky, J.C. Miller, A.B. Robbins, E.J. Rebar, F.D. Urnov, M.C. Holmes, A.M. Pooler, B. Riley, H.S. Zhang, and B.T. Hyman, Persistent repression of tau in the brain using engineered zinc finger protein transcription factors. Sci Adv, 2021. 7(12). PubMed PMID: 33741591; PMCID: PMC7978433.

20. Bi, M., A. Gladbach, J. van Eersel, A. Ittner, M. Przybyla, A. van Hummel, S.W. Chua, J. van der Hoven, W.S. Lee, J. Müller, J. Parmar, G.v. Jonquieres, H. Stefen, E. Guccione, T. Fath, G.D. Housley, M. Klugmann, Y.D. Ke, and L.M. Ittner, Tau exacerbates excitotoxic brain damage in an animal model of stroke. Nature Communications, 2017. 8(1): p. 473.

21. Cloyd, R.A., J. Koren, 3rd, J.F. Abisambra, and B.N. Smith, Effects of altered tau expression on dentate granule cell excitability in mice. Exp Neurol, 2021. 343: p. 113766. PubMed PMID: 34029610; PMCID: PMC8286323.

22. Gheyara, A.L., R. Ponnusamy, B. Djukic, R.J. Craft, K. Ho, W. Guo, M.M. Finucane, P.E. Sanchez, and L. Mucke, Tau reduction prevents disease in a mouse model of Dravet syndrome. Annals of Neurology, 2014. 76(3): p. 443–456.

23. Marciniak, E., A. Leboucher, E. Caron, T. Ahmed, A. Tailleux, J. Dumont, T. Issad, E. Gerhardt, P. Pagesy, M. Vileno, C. Bournonville, M. Hamdane, K. Bantubungi, S. Lancel, D. Demeyer, S. Eddarkaoui, E. Vallez, D. Vieau, S. Humez, E. Faivre, B. Grenier-Boley, T.F. Outeiro, B. Staels, P. Amouyel, D. Balschun, L. Buee, and D. Blum, Tau deletion promotes brain insulin resistance. Journal of Experimental Medicine, 2017. 214(8): p. 2257–2269.

24. Morris, M., P. Hamto, A. Adame, N. Devidze, E. Masliah, and L. Mucke, Age-appropriate cognition and subtle dopamine-independent motor deficits in aged tau knockout mice. Neurobiol Aging, 2013. 34(6): p. 1523–9. PubMed PMID: 23332171; PMCID: 3596503.

25. Cantero, J.L., E. Hita-Yañez, B. Moreno-Lopez, F. Portillo, A. Rubio, and J. Avila, Tau Protein Role in Sleep-Wake Cycle. Journal of Alzheimer’s Disease, 2010. 21: p. 411–421.

26. Fuster-Matanzo, A., M. Llorens-Martín, J. Jurado-Arjona, J. Avila, and F. Hernández, Tau Protein and Adult Hippocampal Neurogenesis. Frontiers in Neuroscience, 2012. 6(104).

27. Ma, Q.-L., X. Zuo, F. Yang, O.J. Ubeda, D.J. Gant, M. Alaverdyan, N.C. Kiosea, S. Nazari, P.P. Chen, F. Nothias, P. Chan, E. Teng, S.A. Frautschy, and G.M. Cole, Loss of MAP Function Leads to Hippocampal Synapse Loss and Deficits in the Morris Water Maze with Aging. The Journal of Neuroscience, 2014. 34(21): p. 7124–7136.

28. Lei, P., S. Ayton, D.I. Finkelstein, L. Spoerri, G.D. Ciccotosto, D.K. Wright, B.X.W. Wong, P.A. Adlard, R.A. Cherny, L.Q. Lam, B.R. Roberts, I. Volitakis, G.F. Egan, C.A. McLean, R. Cappai, J.A. Duce, and A.I. Bush, Tau deficiency induces parkinsonism with dementia by impairing APP-mediated iron export. Nature Medicine, 2012. 18(2): p. 291–295.

29. Li, Z., A.M. Hall, M. Kelinske, and E.D. Roberson, Seizure resistance without parkinsonism in aged mice after tau reduction. Neurobiology of Aging, 2014. 35(11): p. 2617–2624.

30. Chen, R.P., A.S. Gaynor, and W. Chen, Synthetic biology approaches for targeted protein degradation. Biotechnology Advances, 2019. 37(8): p. 107446.

31. Zhao, L., J. Zhao, K. Zhong, A. Tong, and D. Jia, Targeted protein degradation: mechanisms, strategies and application. Signal Transduction and Targeted Therapy, 2022. 7(1): p. 113.

32. Wu, T., H. Yoon, Y. Xiong, S.E. Dixon-Clarke, R.P. Nowak, and E.S. Fischer, Targeted protein degradation as a powerful research tool in basic biology and drug target discovery. Nature Structural & Molecular Biology, 2020. 27(7): p. 605–614.

33. Garber, K., The PROTAC gold rush. Nature Biotechnology, 2022. 40(1): p. 12–16.

34. Martinez-Vicente, M., G. Sovak, and A.M. Cuervo, Protein degradation and aging. Experimental Gerontology, 2005. 40(8): p. 622–633.

35. Basisty, N., J.G. Meyer, and B. Schilling, Protein Turnover in Aging and Longevity. PROTEOMICS, 2018. 18(5-6): p. 1700108.

36. Shringarpure, R. and K.J.A. Davies, Protein turnover by the proteasome in aging and disease1, 2 1This article is part of a series of reviews on “Oxidatively Modified Proteins in Aging and Disease.” The full list of papers may be found on the homepage of the journal.Davies and Shringarpure are studying the mechanism by which the proteasome recognizes and degrades oxidatively damaged proteins, and how protein oxidation and proteolysis are affected by aging and disease. 2Guest Editor: Earl Stadtman. Free Radical Biology and Medicine, 2002. 32(11): p. 1084–1089.

37. Hipp, M.S., S.-H. Park, and F.U. Hartl, Proteostasis impairment in protein-misfolding and -aggregation diseases. Trends in Cell Biology, 2014. 24(9): p. 506–514.

38. Peters, J.M., W.W. Franke, and J.A. Kleinschmidt, Distinct 19 S and 20 S subcomplexes of the 26 S proteasome and their distribution in the nucleus and the cytoplasm. J Biol Chem, 1994. 269(10): p. 7709–18. PubMed PMID: 8125997.

39. Breusing, N. and T. Grune, Regulation of proteasome-mediated protein degradation during oxidative stress and aging. Biol Chem, 2008. 389(3): p. 203–9. PubMed PMID: 18208355.

40. Voss, P. and T. Grune, The nuclear proteasome and the degradation of oxidatively damaged proteins. Amino Acids, 2007. 32(4): p. 527–34. PubMed PMID: 17103119.

41. Bader, N., T. Jung, and T. Grune, The proteasome and its role in nuclear protein maintenance. Exp Gerontol, 2007. 42(9): p. 864–70. PubMed PMID: 17532163.

42. Merker, K., O. Ullrich, H. Schmidt, N. Sitte, and T. Grune, Stability of the nuclear protein turnover during cellular senescence of human fibroblasts. Faseb j, 2003. 17(13): p. 1963–5. PubMed PMID: 12897070.

43. Li, G., W. Ci, S. Karmakar, K. Chen, R. Dhar, Z. Fan, Z. Guo, J. Zhang, Y. Ke, L. Wang, M. Zhuang, S. Hu, X. Li, L. Zhou, X. Li, Matthew F. Calabrese, Edmond R. Watson, Sandip M. Prasad, C. Rinker-Schaeffer, Scott E. Eggener, T. Stricker, Y. Tian, Brenda A. Schulman, J. Liu, and Kevin P. White, SPOP Promotes Tumorigenesis by Acting as a Key Regulatory Hub in Kidney Cancer. Cancer Cell, 2014. 25(4): p. 455–468.

44. Usher, E.T., N. Sabri, R. Rohac, A.K. Boal, T. Mittag, and S.A. Showalter, Intrinsically disordered substrates dictate SPOP subnuclear localization and ubiquitination activity. Journal of Biological Chemistry, 2021. 296: p. 100693.

45. Shin, Y.J., S.K. Park, Y.J. Jung, Y.N. Kim, K.S. Kim, O.K. Park, S.H. Kwon, S.H. Jeon, A. Trinh le, S.E. Fraser, Y. Kee, and B.J. Hwang, Nanobody-targeted E3-ubiquitin ligase complex degrades nuclear proteins. Sci Rep, 2015. 5: p. 14269. PubMed PMID: 26373678; PMCID: PMC4571616.

46. Yamaguchi, N., T. Colak-Champollion, and H. Knaut, zGrad is a nanobody-based degron system that inactivates proteins in zebrafish. Elife, 2019. 8. PubMed PMID: 30735119; PMCID: PMC6384026.

47. Zhuang, M., M.F. Calabrese, J. Liu, M.B. Waddell, A. Nourse, M. Hammel, D.J. Miller, H. Walden, D.M. Duda, S.N. Seyedin, T. Hoggard, J.W. Harper, K.P. White, and B.A. Schulman, Structures of SPOP-substrate complexes: insights into molecular architectures of BTB-Cul3 ubiquitin ligases. Mol Cell, 2009. 36(1): p. 39–50. PubMed PMID: 19818708; PMCID: PMC2847577.

48. Breeuwer, M. and D.S. Goldfarb, Facilitated nuclear transport of histone H1 and other small nucleophilic proteins. Cell, 1990. 60(6): p. 999–1008.

49. Boyden, E.S., F. Zhang, E. Bamberg, G. Nagel, and K. Deisseroth, Millisecond-timescale, genetically targeted optical control of neural activity. Nat Neurosci, 2005. 8(9): p. 1263–8. PubMed PMID: 16116447.

50. Mayford, M., M.E. Bach, Y.-Y. Huang, L. Wang, R.D. Hawkins, and E.R. Kandel, Control of memory formation through regulated expression of a CaMKII transgene. Science, 1996. 274(5293): p. 1678–1683.

51. Cho, Y.K., D. Park, A. Yang, F. Chen, A.S. Chuong, N.C. Klapoetke, and E.S. Boyden, Multidimensional screening yields channelrhodopsin variants having improved photocurrent and order-of-magnitude reductions in calcium and proton currents. J Biol Chem, 2019. 294(11): p. 3806–3821. PubMed PMID: 30610117.

52. Li, D. and Y.K. Cho, High specificity of widely used phospho-tau antibodies validated using a quantitative whole-cell based assay. J Neurochem, 2020. 152(1): p. 122–135. PubMed PMID: 31325178; PMCID: PMC6928439.

53. Wang, S. and Y.K. Cho, An Optimized Calcium-Phosphate Transfection Method for Characterizing Genetically Encoded Tools in Primary Neurons. Methods Mol Biol, 2016. 1408: p. 243–9. PubMed PMID: 26965127.

54. Otsu, N., A Threshold Selection Method from Gray-Level Histograms. IEEE Transactions on Systems, Man, and Cybernetics, 1979. 9(1): p. 62–66.

55. Nagai, Y., T. Kojima, Y. Muro, T. Hachiya, Y. Nishizawa, T. Wakabayashi, and M. Hagiwara, Identification of a novel nuclear speckle-type protein, SPOP. FEBS Letters, 1997. 418(1-2): p. 23–26.

56. Errington, W.J., M.Q. Khan, S.A. Bueler, J.L. Rubinstein, A. Chakrabartty, and G.G. Privé, Adaptor protein self-assembly drives the control of a cullin-RING ubiquitin ligase. Structure, 2012. 20(7): p. 1141–53. PubMed PMID: 22632832.

57. Saerens, D., M. Pellis, R. Loris, E. Pardon, M. Dumoulin, A. Matagne, L. Wyns, S. Muyldermans, and K. Conrath, Identification of a universal VHH framework to graft non-canonical antigen-binding loops of camel single-domain antibodies. J Mol Biol, 2005. 352(3): p. 597–607. PubMed PMID: 16095608.

58. Fridy, P.C., Y. Li, S. Keegan, M.K. Thompson, I. Nudelman, J.F. Scheid, M. Oeffinger, M.C. Nussenzweig, D. Fenyo, B.T. Chait, and M.P. Rout, A robust pipeline for rapid production of versatile nanobody repertoires. Nat Methods, 2014. 11(12): p. 1253–60. PubMed PMID: 25362362; PMCID: PMC4272012.

59. Kubala, M.H., O. Kovtun, K. Alexandrov, and B.M. Collins, Structural and thermodynamic analysis of the GFP:GFP-nanobody complex. Protein Sci, 2010. 19(12): p. 2389–401. PubMed PMID: 20945358; PMCID: 3009406.

60. Ray, M., R. Tang, Z. Jiang, and V.M. Rotello, Quantitative tracking of protein trafficking to the nucleus using cytosolic protein delivery by nanoparticle-stabilized nanocapsules. Bioconjug Chem, 2015. 26(6): p. 1004–7. PubMed PMID: 26011555; PMCID: PMC4743495.

61. Kalderon, D., W.D. Richardson, A.F. Markham, and A.E. Smith, Sequence requirements for nuclear location of simian virus 40 large-T antigen. Nature, 1984. 311(5981): p. 33–38.

62. Dang, C.V. and W.M. Lee, Identification of the human c-myc protein nuclear translocation signal. Molecular and Cellular Biology, 1988. 8(10): p. 4048–4054.

63. Dingwall, C., J. Robbins, S.M. Dilworth, B. Roberts, and W.D. Richardson, The nucleoplasmin nuclear location sequence is larger and more complex than that of SV-40 large T antigen. Journal of Cell Biology, 1988. 107(3): p. 841–849.

64. Kaczmarczyk, S.J., K. Sitaraman, T. Hill, J.L. Hartley, and D.K. Chatterjee, Tus, an E. coli protein, contains mammalian nuclear targeting and exporting signals. PLoS One, 2010. 5(1): p. e8889. PubMed PMID: 20126275; PMCID: PMC2811178.

65. Ramsden, M., L. Kotilinek, C. Forster, J. Paulson, E. McGowan, K. SantaCruz, A. Guimaraes, M. Yue, J. Lewis, and G. Carlson, Age-dependent neurofibrillary tangle formation, neuron loss, and memory impairment in a mouse model of human tauopathy (P301L). Journal of Neuroscience, 2005. 25(46): p. 10637–10647.

66. Götz, J., A. Probst, M.G. Spillantini, T. Schäfer, R. Jakes, K. Bürki, and M. Goedert, Somatodendritic localization and hyperphosphorylation of tau protein in transgenic mice expressing the longest human brain tau isoform. The EMBO Journal, 1995. 14(7): p. 1304–1313.

67. Iwata, M., S. Watanabe, A. Yamane, T. Miyasaka, and H. Misonou, Regulatory mechanisms for the axonal localization of tau protein in neurons. Molecular Biology of the Cell, 2019. 30(19): p. 2441–2457. PubMed PMID: 31364926.

68. DeVos, S.L., D.K. Goncharoff, G. Chen, C.S. Kebodeaux, K. Yamada, F.R. Stewart, D.R. Schuler, S.E. Maloney, D.F. Wozniak, F. Rigo, C.F. Bennett, J.R. Cirrito, D.M. Holtzman, and T.M. Miller, Antisense Reduction of Tau in Adult Mice Protects against Seizures. The Journal of Neuroscience, 2013. 33(31): p. 12887–12897.

69. DeVos, S.L., R.L. Miller, K.M. Schoch, B.B. Holmes, C.S. Kebodeaux, A.J. Wegener, G. Chen, T. Shen, H. Tran, B. Nichols, T.A. Zanardi, H.B. Kordasiewicz, E.E. Swayze, C.F. Bennett, M.I. Diamond, and T.M. Miller, Tau reduction prevents neuronal loss and reverses pathological tau deposition and seeding in mice with tauopathy. Science Translational Medicine, 2017. 9(374): p. eaag0481.

70. Xu, H., T.W. Rösler, T. Carlsson, A. de Andrade, O. Fiala, M. Hollerhage, W.H. Oertel, M. Goedert, A. Aigner, and G.U. Höglinger, Tau silencing by siRNA in the P301S mouse model of tauopathy. Curr Gene Ther, 2014. 14(5): p. 343–51. PubMed PMID: 25687501.

71. Silva, M.C., F.M. Ferguson, Q. Cai, K.A. Donovan, G. Nandi, D. Patnaik, T. Zhang, H.-T. Huang, D.E. Lucente, B.C. Dickerson, T.J. Mitchison, E.S. Fischer, N.S. Gray, and S.J. Haggarty, Targeted degradation of aberrant tau in frontotemporal dementia patient-derived neuronal cell models. eLife, 2019. 8: p. e45457.

72. Chu, T.T., N. Gao, Q.Q. Li, P.G. Chen, X.F. Yang, Y.X. Chen, Y.F. Zhao, and Y.M. Li, Specific Knockdown of Endogenous Tau Protein by Peptide-Directed Ubiquitin-Proteasome Degradation. Cell Chem Biol, 2016. 23(4): p. 453–61. PubMed PMID: 27105281.

73. Lu, M., T. Liu, Q. Jiao, J. Ji, M. Tao, Y. Liu, Q. You, and Z. Jiang, Discovery of a Keap1-dependent peptide PROTAC to knockdown Tau by ubiquitination-proteasome degradation pathway. Eur J Med Chem, 2018. 146: p. 251–259. PubMed PMID: 29407955.

74. Gallardo, G., C.H. Wong, S.M. Ricardez, C.N. Mann, K.H. Lin, C.E.G. Leyns, H. Jiang, and D.M. Holtzman, Targeting tauopathy with engineered tau-degrading intrabodies. Mol Neurodegener, 2019. 14(1): p. 38. PubMed PMID: 31640765; PMCID: PMC6805661.

75. Wagner, U., M. Utton, J.M. Gallo, and C.C. Miller, Cellular phosphorylation of tau by GSK-3 beta influences tau binding to microtubules and microtubule organisation. Journal of Cell Science, 1996. 109(6): p. 1537–1543.

76. Busciglio, J., A. Lorenzo, J. Yeh, and B.A. Yankner, β-Amyloid fibrils induce tau phosphorylation and loss of microtubule binding. Neuron, 1995. 14(4): p. 879–888.

77. Cho, J.-H. and G.V.W. Johnson, Glycogen Synthase Kinase 3β Phosphorylates Tau at Both Primed and Unprimed Sites: DIFFERENTIAL IMPACT ON MICROTUBULE BINDING *. Journal of Biological Chemistry, 2003. 278(1): p. 187–193.

78. Sengupta, A., J. Kabat, M. Novak, Q. Wu, I. Grundke-Iqbal, and K. Iqbal, Phosphorylation of tau at both Thr 231 and Ser 262 is required for maximal inhibition of its binding to microtubules. Arch Biochem Biophys, 1998. 357(2): p. 299–309. PubMed PMID: 9735171.

79. Lin, Y.T., J.T. Cheng, L.C. Liang, C.Y. Ko, Y.K. Lo, and P.J. Lu, The binding and phosphorylation of Thr231 is critical for Tau’s hyperphosphorylation and functional regulation by glycogen synthase kinase 3beta. J Neurochem, 2007. 103(2): p. 802–13. PubMed PMID: 17680984.

80. Cho, J.H. and G.V. Johnson, Primed phosphorylation of tau at Thr231 by glycogen synthase kinase 3beta (GSK3beta) plays a critical role in regulating tau’s ability to bind and stabilize microtubules. J Neurochem, 2004. 88(2): p. 349–58. PubMed PMID: 14690523.

81. Lasagna-Reeves, C.A., D.L. Castillo-Carranza, U. Sengupta, M.J. Guerrero-Munoz, T. Kiritoshi, V. Neugebauer, G.R. Jackson, and R. Kayed, Alzheimer brain-derived tau oligomers propagate pathology from endogenous tau. Sci Rep, 2012. 2: p. 700. PubMed PMID: 23050084; PMCID: PMC3463004.

82. Eftekharzadeh, B., J.G. Daigle, L.E. Kapinos, A. Coyne, J. Schiantarelli, Y. Carlomagno, C. Cook, S.J. Miller, S. Dujardin, A.S. Amaral, J.C. Grima, R.E. Bennett, K. Tepper, M. DeTure, C.R. Vanderburg, B.T. Corjuc, S.L. DeVos, J.A. Gonzalez, J. Chew, S. Vidensky, F.H. Gage, J. Mertens, J. Troncoso, E. Mandelkow, X. Salvatella, R.Y.H. Lim, L. Petrucelli, S. Wegmann, J.D. Rothstein, and B.T. Hyman, Tau Protein Disrupts Nucleocytoplasmic Transport in Alzheimer’s Disease. Neuron, 2018. 99(5): p. 925–940 e7. PubMed PMID: 30189209; PMCID: PMC6240334.

