## Supplementary Materials for "Targeted degradation of microtubule-associated protein tau using an engineered nanobody-E3 ubiquitin ligase adapter fusion"

**Supplementary Table 1.**
Primers used to construct new plasmids used in this study.

| Plasmid name | Primers | Sequence |
|---|---|---|
| pN3-cAbGFP4-SPOP (cAbGFP4 part) | Forward | GGATCCGCCACCATGGATCAAGTCCAACTGGT |
| | Reverse | ACTAGTAGATCCGCTGGAGACGGTGACC |
| pN3-cAbGFP4-SPOP (SPOP part) | Forward | ACTAGTGTCAACATTTCTGGCCAGAAT |
| | Reverse | TGTACAGTTAGGATTGCTTCAGGCGT |
| pN3-cAbGFP4-SPOP-cMyc | Forward | GGATCCGCCACCATGGATCAAGTCCAACTGGT |
| | Reverse | TGTACAGTTAATCCAGCTTCACTCTCTTGGCGGCAGGGCGTGGGGGTCCCAG |
| pN3-cAbGFP4-SPOP-SV40 | Forward | GGATCCGCCACCATGGATCAAGTCCAACTGGT |
| | Reverse | CTGGGACCCCCACGCCCCAAGAAAAAGCGGAAGGTGTAACTGTACA |
| pN3-cAbGFP4-SPOP-NLP | Forward | GGATCCGCCACCATGGATCAAGTCCAACTGGT |
| | Reverse | TGTACAGTTAGTCCAGCTTCTTCTTCTTGGCCTGGCCGGCTTTCTTTGTGGCGGCAGGTCTTTTCACGGCGCGTGGGGGTCCCAG |
| pN3-cAbGFP4-SPOP-Tus | Forward | GGATCCGCCACCATGGATCAAGTCCAACTGGT |
| | Reverse | TGTACAGTTACTTCACGGGCCGCTTGATCTTCAGCTTGCGTGGGGGTCCCAG |
| pN3-cAbGFP4-SPOPΔNLS | Forward | GGATCCGCCACCATGGATCAAGTCCAACTGGT |
| | Reverse | TGTACAGTTAGCGTGGGGGTCCCAG |
| pN3-LaG2-SPOP | Forward | GGATCCGCCACCATGGCTCAGGTGCAGC |
| | Reverse | ACTAGTAGATCCCACGGTCACTTGGGTG |
| pN3-LaM4-SPOP | Forward | GGATCCGCCACCATGGCTCAGGTGCAG |
| | Reverse | ACTAGTAGATCCGGTGAAGGGGCTGGA |
| pN3-LaM4-SPOPΔNLS | Forward | GGATCCGCCACCATGGCTCAGGTGCAG |
| | Reverse | TGTACAGTTAGCGTGGGGGTCCCAG |
| Fck EGFP-Tau | Forward 1 | GGATCCGCCACCATGGTGAGCAAGGGCGAGG |
| | Reverse 1 | GCGGGGCTCAGCCATGGACCCCTTGTACAGCTCGTCCATG |
| | Forward 2 | CATGGACGAGCTGTACAAGGGGTCCATGGCTGAGCCCCGC |
| | Reverse 2 | GAATTCTCACAAACCCTGCTTGGC |
| Fck cAbGFP4-SPOP | Forward | AAGCTTCGAATTCTGC |
| | Reverse | TAATGCTAGCTTAGGATTGCTTCAGGCG |
| Fck cAbGFP4-SPOP-Tus | Forward | AAGCTTCGAATTCTGC |
| | Reverse | TAATGCTAGCTTACTTCACGGGCCGC |
| Fck cAbGFP4-SPOPΔNLS | Forward | AAGCTTCGAATTCTGC |
| | Reverse | TAATGCTAGCTTAGCGTGGGGGTCC |
